# ABioTrans: A Biostatistical tool for Transcriptomics Analysis

**DOI:** 10.1101/616300

**Authors:** Zou Yutong, Bui Thuy Tien, Kumar Selvarajoo

**Author notes:** Equal contribution (Zou Yutong, Bui Thuy Tien). Correspondence to KS.

## Abstract

Here we report a bio-statistical/informatics tool, ABioTrans, developed in R for gene expression analysis. The tool allows the user to directly read RNA-Seq data files deposited in the Gene Expression Omnibus or GEO database. Operated using any web browser application, ABioTrans provides easy options for multiple statistical distribution fitting, Pearson and Spearman rank correlations, PCA, k-means and hierarchical clustering, differential expression analysis, Shannon entropy and noise (square of coefficient of variation) analyses, as well as Gene ontology classifications.

**Availability and implementation:** ABioTrans is available at https://github.com/buithuytien/ABioTrans

Operating system(s): Platform independent (web browser)

Programming language: R (R studio)

Other requirements: Bioconductor genome wide annotation databases, R-packages (shiny, LSD, fitdistrplus, actuar, entropy, moments, RUVSeq, edgeR, DESeq2, NOISeq, AnnotationDbi, ComplexHeatmap, circlize, clusterProfiler, reshape2, DT, plotly, shinycssloaders, dplyr, ggplot2). These packages will automatically be installed when the ABioTrans.R is executed in R studio.

No restriction of usage for non-academic.

## Introduction

Large-scale gene expression analysis requires specialized statistical or bioinformatics tools to rigorously interpret the complex multi-dimensional data, especially when comparing between genotypes. There are already several such tools developed with fairly user-friendly features [1–3]. Nevertheless, there still is a need for more specialised, focused and “click-and-go” analysis tools for different groups of bioinformaticians and wet biologists. In particular, software tools that perform gene expression variability through entropy and noise analyses are lacking. Here, we focused on very commonly used statistical techniques, namely, Pearson and Spearman rank correlations, Principal Component Analysis (PCA), k-means and hierarchical clustering, Shannon entropy, noise (square of coefficient of variation), differential expression analysis and gene ontology classifications [4–7].

Using R programming as the backbone, we developed a web-browser based user interface to simply perform the above-mentioned analyses by a click of a few buttons, rather than using a command line execution. Our interface is specifically made simple considering wet lab biologists as the main users. Nevertheless, our tool will also benefit bioinformaticians and computational biologists at large, as it saves much time for running the R script files for analyses and saving the results in pdf.

## Main Interface & Data Input

Upon loading ABioTrans.R, the homepage window pops up and displays a panel to choose the RNA-Seq data and supporting files (Fig.1). The data file, in comma-separated value (.csv) format, should contain the gene names in rows and genotypes (conditions: wildtype, mutants, replicates, etc.) in columns, following the usual format of files deposited in the GEO database [8]. Supporting files (if applicable) include gene length, list of negative control genes, and metadata file. If the data files contain raw read counts, the user can perform normalization using 5 popular methods: FPKM, RPKM, TPM, Remove Unwanted Variation (RUV) or upper quartile in the pre-processing step [9–12]. FPKM, RPKM and TPM normalization requires inputting gene length file, which should provide matching gene name and their length in base pair in two-column csv file. RUV normalization requires a list of negative control genes (genes that are stably expressed in all experimental conditions), which should be contained in a one-column csv file. If negative control genes are not available, Upper-quartile normalization option will replace RUV. The metadata file is required for differential expression analysis, and should specify experimental conditions (eg. Control, Treated, etc.) for each genotype listed in the data file. Otherwise, the user can move to the next option to perform/click all available analysis buttons (scatter plot, distribution fit, Pearson Correlation, etc.) once a data file is loaded (whether normalized or in raw count).

**Figure 1.**
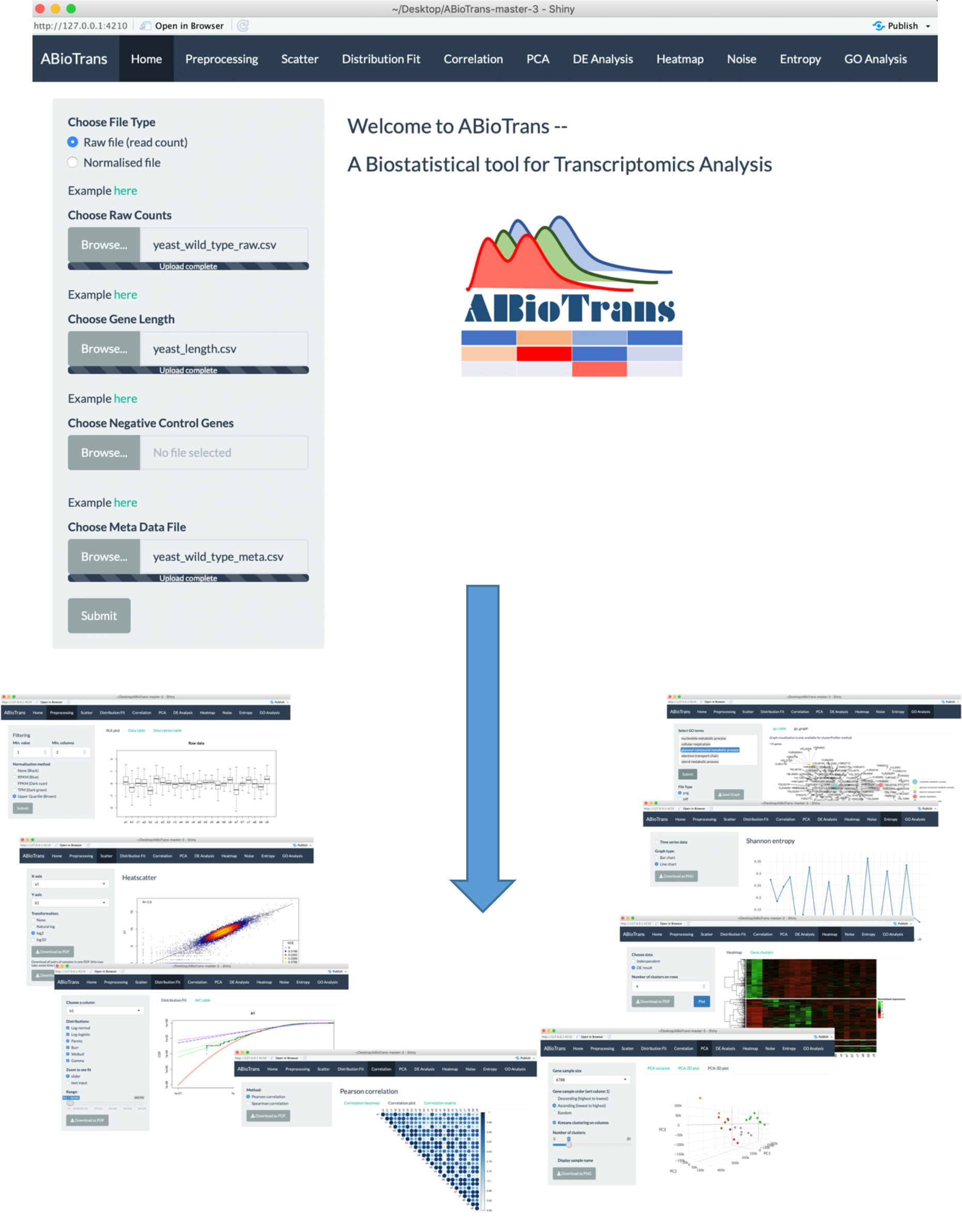
ABioTrans main interface and snapshots of various analysis mode.

## Data Pre-processing

Upon submitting data files and all supporting files (gene length, negative control genes, metadata table), the user can filter the lowly expressed genes by indicating the minimum expression value and the minimum number of samples that are required to exceed the threshold for each gene. If input data contain raw read counts, user can choose one of the normalization options (FPKM, RPKM, TPM, Upper quartile and RUV) listed upon availability of supporting files. FPKM, RPKM and TPM option perform normalization for sequencing depth and gene length, whereas RUV and upper quartile eliminate unwanted variation between samples. To check for sample variation, Relative Log Expression (RLE) plots [13] of input and processed data are displayed for comparison.

## Scatter Plot & Distributions

The scatter plot displays all gene expressions between any two columns selected from the datafile. This is intended to show, transcriptome-wide, how each gene expression varies between any two samples. The lower the scatter, the more similar the global responses and vice-versa [5]. That is, this option allows the user to get an indication of how variable the gene expressions are between any two samples (e.g. between 2 different genotypes or replicates).

After knowing this information, the next process is to make a distribution (cumulative distribution function) plot and compare with the common statistical distributions. As gene expressions are known to follow certain statistical distributions such as power-law or lognormal [14–16], we included the distribution test function. Previously, we have used power-law distribution to perform low signal-to-noise expression cutoff with FPKM expression threshold of less than 10 [7]. Thus, this mode allows the user to check the deviation of their expression pattern with appropriate statistical distributions to select reliable genes for further analysis.

ABioTrans allows the comparison with i) log-normal, ii) Pareto or power-law, iii) log-logistic iv) gamma, v) Weibull and vi) Burr distributions. To compare the quality of statistical distribution fit, the Akaike information criterion (AIC) can also be evaluated on this screen.

## Pearson and Spearman Correlations

This mode allows the user to compute linear (Pearson) and monotonic non-linear (Spearman) correlations, i) in actual values in a table or ii) as a density gradient plot between the samples.

## PCA and K-means clustering

The PCA button plots the variance of all principal components and allows 2-D and 3-D plots of any PC-axis combination. There is also a slide bar selector for testing the number of k-means clusters.

## Entropy and Noise

These functions measure the disorder or variability between samples using Shannon entropy and expressions scatter [17, 18]. Entropy values are obtained through binning approach and the number of bins are determined using Doane’s rule [5, 19].

To quantify gene expressions scatter, the noise function computes the squared coefficient of variation [13], defined as the variance (σ^2^) of expression divided by the square mean expression (μ^2^), for all genes between all possible pairs of samples [5].

## Differential Expression Analysis

AbioTrans provides users with 3 options to carry out differential expression (DE) analysis on data with replicates: edgeR, DESeq2 and NOISeq [20–22]. In case there are no replicates available for any of the experimental condition, technical replicates can be simulated by NOISeq. edgeR and DESeq2 requires filtered raw read counts, therefore, it is recommended that the user provide input data file containing raw counts if DE analysis is required using either of the two methods. On the other hand, if only normalized gene expression data is available, NOISeq is recommended.

To better visualize DE analysis result by edgeR and DESeq, volcano plot (plot of log_10_-p-value and log_2_-fold change for all genes) distinguishing the significant and insignificant, DE and non-DE genes, is displayed. Plot of dispersion estimation, which correlates to gene variation, is also available in accordance to the selected analysis method.

## Hierarchical clustering & Heatmap

This function allows clustering of differentially expressed genes. User can either utilize the result from DE analysis, or carry out clustering independently by indicating the minimum fold change between 2 genotypes.

For clustering independently, normalized gene expression (output from pre-processing tab) first undergo scaling defined by 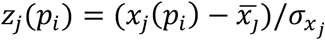 where *z*_*j*_(*p*_*i*_) is the scaled expression of the *j*^*th*^ gene, *x*_*j*_(*p*_*i*_) is expression of the *j*^*th*^ gene in sample *p*_*i*_, 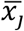 is the mean expression across all samples and 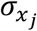 is the standard deviation [23]. Subsequently, Ward hierarchical clustering is applied on the scaled normalized gene expression.

ABioTrans also lists the name of genes for each cluster.

## Gene Ontology

This function is used to define the biological processes or enrichment of differentially regulated genes in a chosen sample or cluster. User can select among 3 gene ontology enrichment test: enrichR, clusterProfiler and GOstats [24–26].

The user needs to create a new csv file providing the name of genes (for each cluster) lining in 1 column (foreground genes). Background genes (or reference genes), if available, should be prepared in the same format. Next, the sample species, gene ID type (following NCBI database [8]) and one of the three subontology (biological process, molecular function or cellular component) need selection. The output results in a gene list, graph (clusterProfiler), and pie chart (clusterProfiler and GOstats) for each ontology.

## Typical Analysis Time Estimation

The loading time ABioTrans for a first time R user is about 30 min on a typical Windows notebook or Macbook. This is due to the installation of the various R-packages that are prerequisite to run ABioTrans. For regular R users, who have installed most packages, the initial loading can take between a few to several minutes depending on whether package updates are required. Once loaded, the subsequent re-load will take only a few seconds.

The typical time taken from pre- to post-processing using all features in ABioTrans is between 10-20 minutes. Table 1 below highlights the typical time taken for each execution for 3 sample data deposited in ABioTrans Github folder (*zfGenes, Biofilm-Yeast, and Yeast-biofilm2*).

**Table 1.** Time comparison of functionalities for different test data.

| Type of Analysis |  | Time (s) |  |  |
| --- | --- | --- | --- | --- |
|  |  | Test 1* | Test 2# | Test 3^ |
| Pre-processing | TPM/RPKM/FPKM and RLE plot | - | - | 0.6 s |
|  | Upper quartile normalization and RLE plot | - | 0.5s | 0.6 s |
|  | RUV normalization and RLE plot | 1.7 s | - | - |
| Scatter Plot |  | 0.01s | 0.01s | 0.01s |
| Distribution fitting (for all 6 distributions) |  | 4.3 s | 3.1s | 2.5s |
| Correlation matrix |  | 0.01 | 0.01s | 0.01s |
| PCA calculation and plotting |  | 0.01 | 0.01 | 0.01 |
| DE analysis | edgeR | 7.89 | 1.52s | 5.23s |
|  | DESeq2 | 15.4 s | 3.1 s | 11.3s |
|  | NOISeq | 29.6 s | 22.87s | 31.0s |
| Heat map and hierarchical clustering | DE (using edgeR result) (5 clusters) | 0.36 s | 1.7 s | 0.25s |
|  | Independent (5 clusters) | 30.4 s | 7.7 s | 4.6s |
| Noise |  | 3.2 s | 1.3 s | 3.9s |
| Shannon Entropy |  | 0.03 s | 0.02 s | 0.08s |
| GO Analysis<br>(using edgeR result) | clusterProlifer | 20.2 s | 10.3 s | 9.1s |
|  | GOstats | 26.6 s | 10.2 s | 12.3s |
|  | EnrichR | - | - | - |

\* ref. 27: GEO accession number: GSE53334, # ref. 28: GEO accession number: GSE85595
^ ref. 29 : GEO accession number: GSE85843

ABioTrans has also been compared with other similar freely available RNA-Seq GUI tools, and it demonstrates better functionalities and capabilities (Supplementary Table S1).

## Summary

ABioTrans is a user-friendly, easy-to-use, point-and-click statistical tool tailored to analyse RNA-Seq datafiles. It can also be used to analyse any highthroughput data as long as they follow the format listed in this technology report.

## Acknowledgement & Funding

The authors thank Lin Yifeng, Ng Shi Yuan and Nic Lindley for discussions, and the Biotransformation Innovation Platform (BioTrans, A*STAR) for funding the work.

## References

1. Poplawski A, Marini F, Hess M, Zeller T, Mazur J, Binder H. Systematically evaluating interfaces for RNA-seq analysis from a life scientist perspective. Brief Bioinform. 2016 Mar;17(2):213–23.

2. Russo F, Angelini C. RNASeqGUI: a GUI for analysing RNA-Seq data. Bioinformatics. 2014 Sep 1;30(17):2514–6.

3. Velmeshev D, Lally P, Magistri M, Faghihi MA. CANEapp: a user-friendly application for automated next generation transcriptomic data analysis. BMC Genomics. 2016 Jan 13;17:49.

4. Tsuchiya M, Piras V, Choi S, Akira S, Tomita M, Giuliani A, Selvarajoo K. Emergent genome-wide control in wildtype and genetically mutated lipopolysaccarides-stimulated macrophages. PLoS One. 2009. 4:e4905

5. Piras V, Tomita M, Selvarajoo K. Transcriptome-wide variability in single embryonic development cells. Sci Rep. 2014. 4:7137.

6. Piras V, Selvarajoo K. The reduction of gene expression variability from single cells to populations follows simple statistical laws. Genomics. 2015. 105:137–144.

7. Simeoni O, Piras V, Tomita M, Selvarajoo K. Tracking global gene expression responses in T cell differentiation. Gene. 2015. 569:259–266.

8. Clough E, Barrett T. The Gene Expression Omnibus Database. Methods Mol Biol. 2016;1418:93–110.

9. Trapnell,C. et al. Transcript assembly and quantification by RNA-seq reveals unannotated transcripts and isoform switching during cell differentiation. Nat. Biotechnol. 2010, 28:511–515.

10. Mortazavi,A. et al. Mapping and quantifying mammalian transcriptomes by RNA-Seq. Nat. Methods 2008, 5:621–628.

11. Wagner GP, Kin K, Lynch VJ. Measurement of mRNA abundance using RNA-seq data: RPKM measure is inconsistent among samples. Theory Biosci. 2012, 131:281–5.

12. Risso D, Ngai J, Speed TP, Dudoit S. Normalization of RNA-seq data using factor analysis of control genes or samples. Nat Biotechnol. 2014, 32:896–902.

13. Gandolfo LC, Speed TP. RLE plots: Visualizing unwanted variation in high dimensional data. PLoS One. 2018, 13:e0191629.

14. Furusawa C, Kaneko K. Zipf’s law in gene expression. Phys Rev Lett. 2003 Feb 28;90(8):088102.

15. Bengtsson M, Stahlberg A, Rorsman P, Kubista M. Gene expression profiling in single cells from the pancreatic islets of Langerhans reveals lognormal distribution of mRNA levels. Genome Res. 2005 Oct;15(10):1388–92.

16. Beal J. Biochemical complexity drives log-normal variation in genetic expression. IET Engineering Biol 2017; 1(1):55–60.

17. Shannon, C. E. A mathematical theory of communication. Bell Syst. Tech. J. 27, 379-423, 623–656 (1948).

18. Bar-Even, A. et al. Noise in protein expression scales with natural protein abundance. Nat. Genet. 38, 636–643 (2006).

19. Doane, D. P. Aesthetic frequency classification. Am. Stat. 30, 181–183 (1976).

20. McCarthy DJ, Chen Y, Smyth GK. Differential expression analysis of multifactor RNA-Seq experiments with respect to biological variation. Nucleic Acids Res. 2012, 40:4288–97.

21. Love MI, Huber W, Anders S. Moderated estimation of fold change and dispersion for RNA-seq data with DESeq2. Genome Biol. 2014, 15:550.

22. Tarazona S, Furió-Tarí P, Turrà D, Pietro AD, Nueda MJ, Ferrer A, Conesa A. Data quality aware analysis of differential expression in RNA-seq with NOISeq R/Bioc package. Nucleic Acids Res. 2015, 43:e140.

23. Simeoni O, Piras V, Tomita M, Selvarajoo K. Tracking global gene expression responses in T cell differentiation. Gene. 2015, 569:259–66.

24. Kuleshov MV et al. Enrichr: a comprehensive gene set enrichment analysis web server 2016 update. Nucleic Acids Res. 2016, 44:W90–7.

25. Yu G, Wang L, Han Y, He Q. clusterProfiler: an R package for comparing biological themes among gene clusters.” OMICS: A Journal of Integrative Biology 2012, 16:284–287.

26. Falcon S, Gentleman R. Using GOstats to test gene lists for GO term association. Bioinformatics. 2007, 23:257–8.

27. Risso D, Ngai J, Speed TP, Dudoit S. Normalization of RNA-seq data using factor analysis of control genes or samples. Nat Biotechnol 2014 Sep;32(9):896–902. PMID:25150836

28. Bendjilali N, et al. Time-Course Analysis of Gene Expression During the Saccharomyces cerevisiae Hypoxic Response. G3 (Bethesda) 2017, 7:221–231.

29. Cromie GA, Tan Z, Hays M, Jeffery EWet al. Dissecting Gene Expression Changes Accompanying a Ploidy-Based Phenotypic Switch. G3 (Bethesda)2017 Jan 5;7(1):233–246. PMID:27836908

