## Supplemental Table 1 for "ABioTrans: A Biostatistical tool for Transcriptomics Analysis"

|  | | | **ABioTrans** | **DEApp** | **DEB** | **DEBrowser** |
| --- | --- | --- | --- | --- | --- | --- |
| Literature reference | | | Zou Y, et al | Li Y & Andrade J [1] | Yao JQ & Yu F [2] | Kucukural A, et al [3] |
| Download link | | | <https://github.com/buithuytien/ABioTrans> | <https://gallery.shinyapps.io/DEApp/> | <http://www.ijbcb.org/DEB/php/onlinetool.php> | <https://github.com/UMMS-Biocore/debrowser> |
| Platform | | | R | Web | Web | R |
| **Analysis:** | | | | | | |
| pre-processing |  | Low count filtering | ✔ | ✔ | NA | ✔ |
|  | Normalization | Sequencing depth correction | TPM, RPKM, FPKM | NA |  | NA |
|  |  | Batch effect correction | UQ, RUV, RLE | unstated method |  | UQ, MRN, RLE |
| PCA | | PCA (2D/ 3D) | 2D & 3D | NA (MDS plot instead) |  | 2D |
|  |  | Scree plot | Scree plot given |  |  | Scree plot given |
| Distribution fitting | | | ✔ | NA |  | NA |
| Scatter plot | | | ✔ |  |  | ✔ |
| Correlation analysis | | | Pearson, spearman |  |  | Spearman |
| DE Analysis | Packages | | edgeR, DESeq2, NOISeq | edgeR, limma, DESeq2 | edgeR, DESeq2, Bayseq | edgeR, limma, DESeq2 |
|  | Supporting plots | | volcano plot, dispersion plot | volcano, dispersion | NA | scatter, volcano, MA plot |
|  | DE cut off criteria | | FDR, FC | FDR, p-value, FC | FDR only | Default |
|  | DE analysis parameters | | Default | Default | Default | User input |
| Heat map | | | ✔ | NA | NA | ✔ |
| Clustering | | | ✔ |  |  | NA |
| Entropy | | | ✔ |  |  |  |
| Noise | | | ✔ |  |  |  |
| Gene ontology | | Packages | ClusterProfiler, GOstats, enrichR |  |  | ClusterProfiler, Disease ontology, compareCluster |
|  |  | Visualization | Pie chart, Graph |  |  | summary plot, dot plot |

1. Li Y, Andrade J. DEApp: an interactive web interface for differential expression analysis of next generation sequence data. Source Code Biol Med. 2017;12. 10.1186/s13029-017-0063-4.

2. Yao JQ, Yu F. DEB: A web interface for RNA-seq digital gene expression analysis. Bioinformation. 2011;7(1):44-5.

3. Alper Kucukural, Onur Yuksel, Deniz M. Ozata, Melissa J. Moore, Manuel Garber, DEBrowser: Interactive Differential Expression Analysis and Visualization Tool for Count Data, BMC Genomics 2019, 20:6 doi: 10.1186/s12864-018-5362-x
